# Accumulation of storage proteins in plant seeds is mediated by amyloid formation

**DOI:** 10.1101/825091

**Authors:** Kirill S. Antonets, Mikhail V. Belousov, Anna I. Sulatskaya, Maria E. Belousova, Anastasiia O. Kosolapova, Maksim I. Sulatsky, Elena A. Andreeva, Pavel A. Zykin, Yury V. Malovichko, Oksana Y. Shtark, Anna N. Lykholay, Kirill V. Volkov, Irina M. Kuznetsova, Konstantin K. Turoverov, Elena Y. Kochetkova, Oleg. N. Demidov, Igor A. Tikhonovich, Anton A. Nizhnikov

## Abstract

Amyloids are protein aggregates with a highly ordered spatial structure giving them unique physicochemical properties. Different amyloids not only participate in the development of numerous incurable diseases but control vital functions in Archaea, Bacteria and Eukarya. Plants are a poorly studied systematic group in the field of amyloid biology. Amyloid properties have not yet been demonstrated for plant proteins under native conditions *in vivo.* Here we show that seeds of garden pea *Pisum sativum* L. contain amyloid-like aggregates of storage proteins, the most abundant one, 7S globulin Vicilin, forms *bona fide* amyloids *in vivo* and *in vitro*. The Vicilin amyloid accumulation increases during seed maturation and wanes at germination. Amyloids of Vicilin resist digestion by gastrointestinal enzymes, persist in canned peas and exhibit toxicity for yeast and mammalian cells. Our finding for the first time reveals involvement of amyloid formation in the accumulation of storage proteins in plant seeds.

## INTRODUCTION

Amyloids represent protein aggregates having an unusual structure formed by intermolecular beta-sheets and stabilized by numerous hydrogen bonds^1^. Such a structure called ‘cross-β’^2^ gives amyloids the morphology of predominantly unbranched fibrils and unique physicochemical properties including (i) resistance to treatment with ionic detergents and proteinases; (ii) binding amyloid-specific dyes like Thioflavin T (ThT); (iii) apple-green birefringence in polarized light upon binding with Congo Red (CR) dye^1,3^.

The biological significance of amyloids is based on two aspects: pathological and functional. Amyloid deposition leads to the development of more than 40 incurable human and animal diseases including various types of amyloidoses and neurodegenerative disorders^4,5^. Nevertheless, amyloids may not be only pathogenic but also functional^6^. A growing number of studies demonstrate that amyloids play vital roles in Archaea^7^, Bacteria and Eukarya including humans^8^. Amyloids of prokaryotes fulfill mostly structural (biofilm and sheaths formation) and storage (toxin accumulation) functions^9^. In fungi, infectious amyloids called prions control heterokaryon incompatibility, multicellularity and drug resistance^10–12^. In animals, amyloid formation is important for different functions including the long-term memory potentiation, melanin polymerization, hormone storage, and programmed necrosis^13^. Compared to other groups of organisms, plants remain to be poorly studied in the field of amyloid biology. However, several plant proteins or their regions were shown to form fibrils with several properties of amyloids *in vitro* (after the proteolytic digestion or other treatments^14,15^) suggesting that plants might form *bona fide* amyloids *in vivo^16^*.

Previously, we performed a large-scale bioinformatic analysis of potentially amyloidogenic properties of plant proteins including all annotated proteomes of land plant species^17^. This screening demonstrated that seed storage proteins comprising the evolutionary conservative β-barrel domain Cupin-1 were rich in amyloidogenic regions in the majority of analyzed species^17^. Such proteins belonging mainly to 11S and 7S globulins^18^ represent key amino acid sources for the growing seedlings, important components of human diet and major allergens^19^. We hypothesize that the amyloid formation could occur at seed maturation to stabilize storage proteins, thus preventing their degradation and misfolding during the seed dormancy. In order to test this hypothesis, we have analyzed whether amyloid proteins are present in seeds of an important agricultural crop and Mendel’s genetic model, garden pea *Pisum sativum* L.

## RESULTS

### The storage proteins form aggregates resistant to treatment with ionic detergents in pea seeds

Resistance to treatment with ionic detergents is a typical feature of amyloids discriminating them from the majority of other non-amyloid protein complexes^20^. We decided to study whether storage proteins form detergent resistant aggregates in plant seeds. For this purpose, we have used mature seeds of garden pea *P. sativum* L. genetic line *Sprint-2* collected at 30 days after the pollination when the accumulation of storage proteins is expected to reach its maximum^21^. We have extracted protein complexes resistant to treatment with ionic detergent sodium dodecyl sulfate (SDS, 1 %) from pea seeds using a previously published method called PSIA-LC-MALDI^22^ with several modifications (see Materials and Methods). Next, the obtained detergent-resistant protein fractions have been solubilized with formic acid and subjected to trypsinolysis followed by the reversed-phase high performance liquid chromatography and mass-spectrometry (see Materials and Methods). As a result, we have identified all three major classes of seed storage proteins (Vicilins, Convicilins and Legumins) and several other proteins including heteropolymeric iron-binding ferritin, biotin-containing protein SBP65 and drought stress response protein dehydrin as detergent resistant components of pea seeds (Table S1). Among identified proteins, 7S storage globulins Vicilins have demonstrated the highest mass-spectrometric scores suggesting for their prevalence in detergent resistant fraction of pea seeds (Table S1). We have selected 47 kDa Vicilin for a detailed analysis of its amyloid properties *in vivo* and *in vitro* since its N- and C-terminal regions have been identified in MS/MS analysis indicating that full-length protein participates in the formation of detergent resistant polymers (Figure S1).

### Recombinant Vicilin and its domains, Cupin-1.1 and Cupin-1.2, form detergent-resistant fibrils *in vitro*

To check ability of Vicilin to aggregate *in vitro*, we produced the full-length C-terminally 6x-His tagged Vicilin in the *E. coli* cells, extracted and purified it. Since Vicilin contains two evolutionary conserved Cupin-1 domains called Cupin-1.1 and Cupin-1.2 (Figure 1a) belonging to the Cupin β-barrel domain superfamily, which have been bioinformatically predicted to be amyloidogenic^17^, we have also analyzed the aggregation of these domains *in vitro* to estimate their contributions to the aggregation of the full-length Vicilin. We have found that incubation of Vicilin, Cupin-1.1 and Cupin-1.2 proteins in phosphate buffer (pH 7.4) for one day at room temperature (25°C) with the constant stirring has caused the formation of typical amyloid fibrils only in the Cupin-1.2 sample. At the same time, Vicilin and Cupin-1.1 have formed mainly morphologically unstructured aggregates visualized by the transmission electron microscopy (TEM) (Figure 1b, top row). Prolonged incubation (two weeks) has caused the formation of more structured Cupin-1.1 but not Vicilin aggregates (Figure 1b, second row). The elevated temperature (50°C) has led to the formation of more compact Vicilin aggregates (Figure 1.b, third row). The best results have been obtained by the usage of Hexafluoro-2-propanol (HFIP) solvent at 37°C that has induced the formation of linear fibrils in all three samples. While Vicilin has only been partially presented by fibrils and contained major fraction of less structured aggregates, almost all aggregates of Cupin-1.1 and Cupin-1.2 in such conditions have exhibited fibrillary morphology typical for amyloids (Figure 1b, bottom row). We have analyzed the resistance of the *in vitro* obtained Vicilin, Cupin-1.1 and Cupin-1.2 aggregates to treatment with ionic detergents. For this purpose, protein samples have been treated for 5 min with the SDS-PAGE sample buffer containing 2% SDS or boiled for 5 min with the same buffer containing 2% SDS and then loaded onto the gel and subjected to the western-blot analysis. The results of this experiment have demonstrated that all three proteins form aggregates resistant to the treatment with both cold and hot SDS with the part of aggregates dissolved after boiling (Figure 1c).

**Figure 1.**
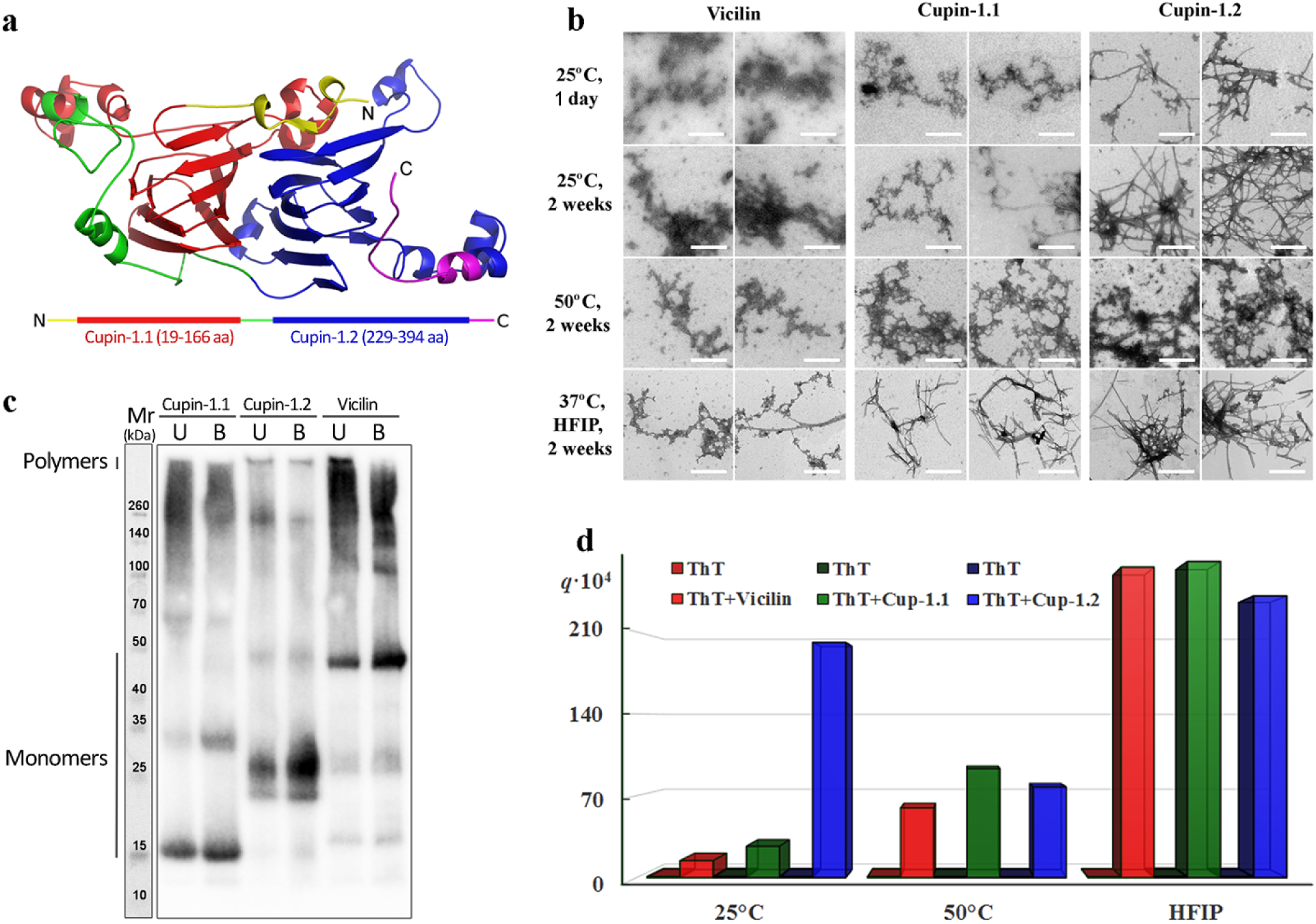
Full-length Vicilin and its domains, Cupin-1.1 and Cupin-1.2, form aggregates with fibrillary morphology that possess detergent resistance and bind amyloid-specific dye ThT. **(a)** The structure of the Vicilin monomer has been predicted by the I-TASSER server^27^. The domain structure of Vicilin is presented below with bars. The Cupin-1.1 domain is denoted by red color and Cupin-1.2 is denoted with blue. The linkers are colored with green and yellow and purple colors. N- and C-termini are shown. **(b)** Transmission electron microscopy (TEM) of the Vicilin, Cupin-1.1 and Cupin-1.2 aggregates obtained in various conditions as indicated. The scale bars correspond to 500 nm. **(c)** The resistance of the Vicilin, Cupin-1.1 and Cupin-1.2 aggregates to the treatment with cold (U, unboiled) and hot (B, boiled) 2 % SDS. Respective molecular weights (kDa) are shown. **(d)** Fluorescence quantum yield of amyloid-specific probe ThT in a free state in buffer solution (left bars) and bound to Vicilin, Cupin-1.1 and Cupin-1.2 aggregates (right bars) have been determined by using the equilibrium microdialysis for the sample preparation.

To confirm the assumption of the Vicilin, Cupin-1.1 and Cupin-1.2 fibril formation we have studied the interaction of the obtained aggregates with ThT^23^. A unique feature of this dye is a very weak fluorescence in the free state in an aqueous solution and an intense fluorescence in the bound to fibrils state. Since the samples contained both ones bound to fibrils and free ThT, we have used an equilibrium microdialysis for preparing test solutions and the separation of photophysical characteristics of two different dye fractions^24^. The use of this approach and other special techniques taking into account the contribution of the aggregates light scattering to the recorded absorption spectra^24^ and the correction of the recorded fluorescence intensity to the primary inner filter effect^25^, have made it possible to calculate the fluorescence quantum yield of free ThT in the samples (the value of which turned out to be close to 10^−4^, which suits the literature data^26^) and the dye associated with the studied aggregates (Figure 1d).

We have found that ThT binds to all the tested samples, however, an increase in the fluorescence quantum yield of the bound dye in comparison to that for free ThT varies considerably in different samples (Figure 1d). The greatest increase in the fluorescence quantum yield is observed in the case of ‘typical’ Cupin-1.2 fibrillar aggregates obtained in the phosphate buffer at 25°C (more than 2 orders of magnitude). At the same time, ThT fluoresces significantly less when it is binding to Vicilin and Cupin-1.1 aggregates obtained in the same conditions (Figure 1d). It can be assumed that the main fraction of the bound dye in these samples interacts with aggregates (apparently with less compact and ordered compared to fibrillar ones) not specifically, which leads only to an insignificant restriction of the intramolecular mobility of ThT fragments relative to one another, and, therefore, to a small increase in its fluorescence quantum yield. ThT bound to Vicilin, Cupin-1.1 and Cupin-1.2 aggregates obtained in the phosphate buffer at 50°C has exhibited the similar intermediate means of the fluorescence quantum yields (Figure 1d) indicating their partially structured morphology (Figure 1b, third row). Finally, ThT bound to Vicilin, Cupin-1.1 and Cupin-1.2 fibrils obtained in HFIP at 37°C has exhibited high fluorescence quantum yields (Figure 1d) confirming their highly structured spatial characteristics.

### Fibrils of the full-length Vicilin, Cupin-1.1 and Cupin-1.2 possess amyloid properties

One of the important structural features of amyloids is their richness in β-sheets^2^. Far-UV circular dichroism (CD) spectra of Cupin-1.1 and Cupin-1.2 aggregates, which are mainly fibrillar (Figure 1b), have apparent negative shoulder around 220 nm^28^ (Figure 2a), that allows us to suggest the essential content of beta-sheets in their structure. At the same time in CD spectrum of Vicilin aggregates this shoulder is significantly smaller and the shoulder around 200 nm (that is typical of unfolded proteins^29^) is predominant (Figure 2a). This does not contradict the results of TEM, indicating a lower content of ordered fibrillar structures in Vicilin sample compared with Cupin-1.1 and Cupin-1.2 samples (Figure 1b). Despite the differences in the recorded CD spectra their quantitative analysis has suggested the high content of ordered beta-structure for all tested aggregates (about 39-41%).

**Figure 2.**
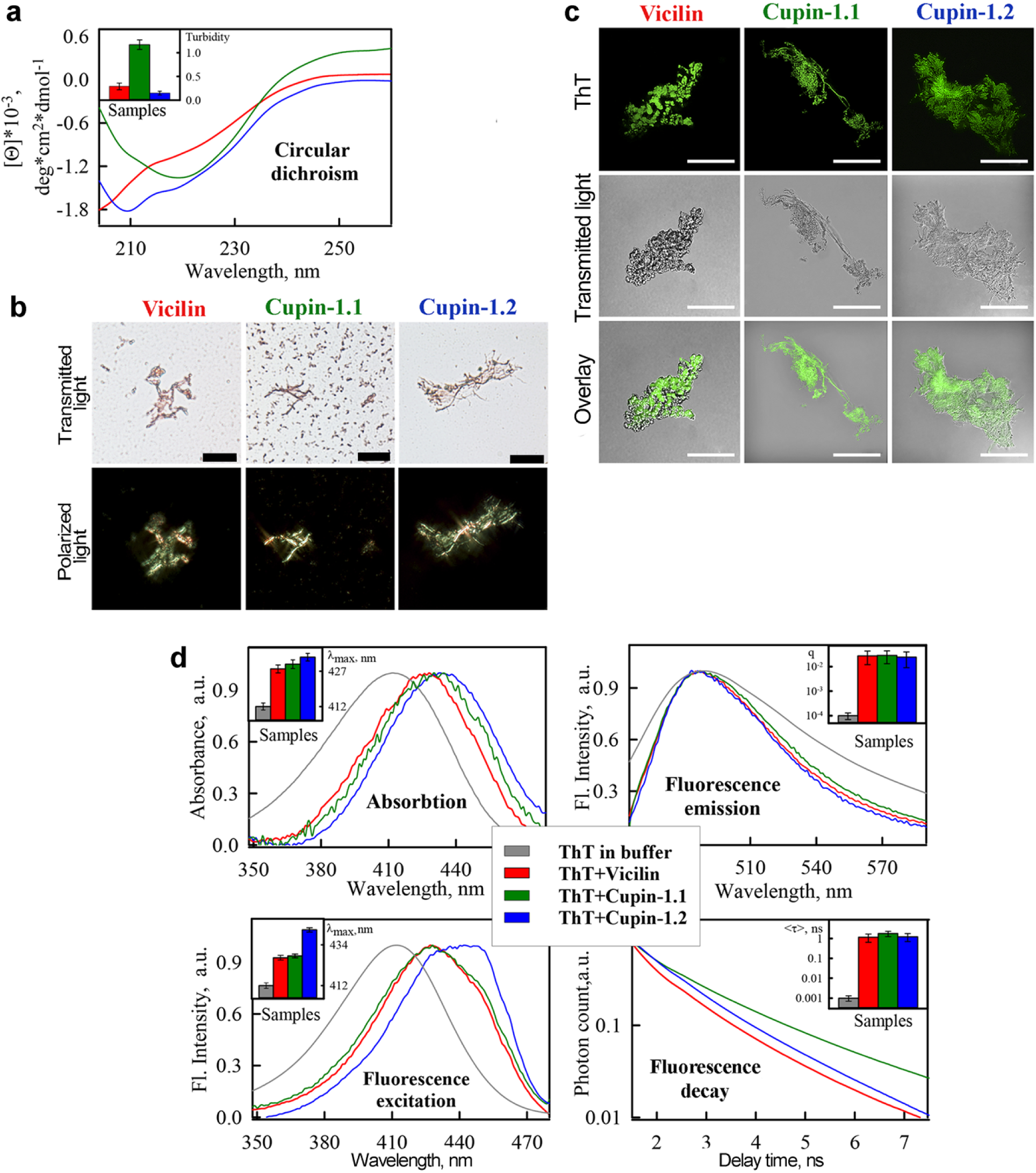
Vicilin, Cupin-1.1 and Cupin-1.2 aggregates exhibit amyloid properties. **(a)** Circular dichroism analysis. The (Inset) shows the turbidity of the samples. **(b)** Polarization microscopy of the Vicilin, Cupin-1.1 and Cupin-1.2 aggregates stained with Congo Red. The scale bar is equal to 20 μm. Top row – transmitted light, Bottom – polarized light. **(c)** Confocal microscopy of the Vicilin, Cupin-1.1 and Cupin-1.2 aggregates stained with ThT. Fluorescence images of the ThT stained aggregates (top row), transmitted light images showing the presence of aggregates in the sample (middle) and the overlay of the images (bottom) is presented. The scale bar is equal to 15 μm. **(d)** Absorption, fluorescence excitation and emission spectra and fluorescence decay curves of the ThT bound to Vicilin, Cupin-1.1 and Cupin-1.2 aggregates. The insets of the corresponding panels show the position of the maxima of the absorption and fluorescence excitation spectra, the values of the fluorescence quantum yield and lifetime of the bound to fibrils dye. Decoding used colors is given in the Figure.

Next, we have stained Vicilin, Cupin-1.1 and Cupin-1.2 aggregates with Congo red dye. When Congo red binds to amyloids it leads to the so-called ‘apple-green birefringence’ in polarized light^30^. This effect is highly specific and for a long time has considered to be the ‘gold standard’ in the amyloid diagnostics^4^. The results of this test have demonstrated that Vicilin, Cupin-1.1 and Cupin-1.2 aggregates bind Congo red and exhibit birefringence (Figure 2b). Notably, despite the fact that Vicilin aggregates obtained in the aforementioned experiments do not exhibit clear fibrillary morphology (Figure 1b), they contain a significant admixture of the amyloid fibrils exhibiting birefringence that is easily detected by the polarization microscopy (Figure 2b).

The amyloid nature of the studied samples has also been proven by their confocal microscopy in the presence of ThT. The dye has stained not only Cupin-1.1 and Cupin-1.2 fibrils, but also less ordered Vicilin aggregates (Figure 2c). Using solutions prepared by the equilibrium microdialysis, we have determined the absorption, fluorescence and fluorescence excitation spectra, as well as the fluorescence decay curves of ThT bound with these aggregates (Figure 2d). We have shown the long-wavelength shift of the dye absorption and fluorescence excitation spectra when it binds to protein aggregates (Figure 2d, Insets of left panels). The determined excitation spectra have a maximum at a wavelength of about 430 nm and longer-wavelength shoulder of about 450 nm, which indicates the existence of two different types of the dye-amyloid binding (Figure 2d, left panel). A binding mode with a maximum of the absorption and fluorescence excitation spectra of about 450 nm was previously detected when ThT binds to amyloid fibrils were formed from insulin and lysozyme^24,31^. Another binding type can be due to the fraction of the dye molecules associated with the less ordered and compact aggregates in the samples. The maximum of fluorescence spectra of ThT bound to Vicilin, Cupin-1.1 and Cupin-1.2 aggregates (Figure 2d, right panel) coincides with the maximum of that of the free dye, as in the case of ThT interaction with amyloid fibrils formed from other amyloidogenic proteins^32^. In addition, we have shown a significant increase in the fluorescence quantum yield and lifetime of the dye bound to tested aggregates (Figure 2d, right panels), which is also a characteristic feature of the ThT interaction with amyloid aggregates.

Thus, based on the wide array of experimental data we may conclude that the full-length Vicilin and both its domains Cupin-1.1 and Cupin-1.2 form amyloid aggregates *in vitro.* The fibril formation by Cupin-1.1 and Cupin-1.2 suggests their contribution to the amyloid formation by the full-length Vicilin. In contrast to both Cupin domains, Vicilin tends to form mixture of *bona fide* amyloid fibrils with less structured aggregates.

### Fibrillogenesis of the full-length Vicilin can be efficiently seeded by the pre-×incubated Vicilin or Cupin-1.2 fibrils

Characteristic feature of many amyloids is their ability to ‘seed’ the aggregation of monomers with small quantities of fibrils^33^. The process of such ‘seeding’ is mostly sequence-specific. Therefore, we have decided to investigate the ability of the Vicilin, Cupin-1.1 and Cupin-1.2 aggregates to induce the fibrillation of the full-length Vicilin. For this purpose, we have used ‘seeds’ prepared from pre-incubated Vicilin, Cupin-1.1 or Cupin-1.2 aggregates. It has turned out that Vicilin aggregates obtained by the usage of the seeds on the basis of Cupin-1.1 aggregates do not differ in their morphology from those prepared in the absence of ‘seeds’ (Figure 3a, middle panel). At the same time, using the ‘seeds’ from the Cupin-1.2 fibrils has resulted in more efficient formation of the Vicilin fibrils (Figure 3a, right panel). Finally, ‘seeds’ from the Vicilin aggregates obtained in previous experiments have caused the formation of almost exclusively Vicilin fibrils with the visual absence of unstructured aggregates (Figure 3a, left panel).

**Figure 3.**
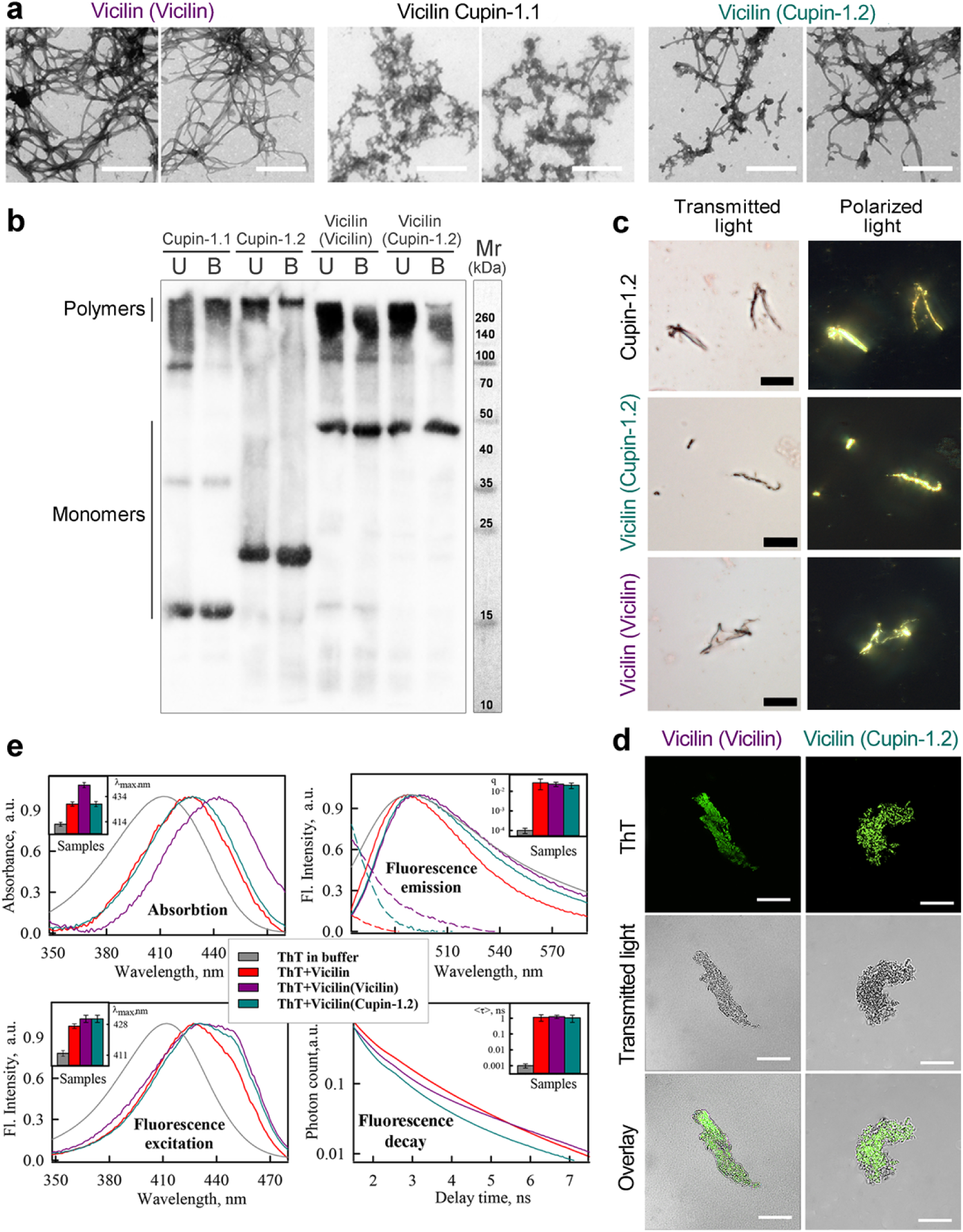
Vicilin amyloid fibril formation is efficiently seeded by its aggregates or Cupin-1.2 fibrils. Seeds used for each of the samples are indicated in parentheses. **(a)** TEM of the Vicilin fibrils obtained by seeding. The scale bar is equal to 500 nm. **(b)** Detergent resistance of the fibrils to cold (U, unboiled) and hot (B, boiled) 2% SDS. Respective molecular weights (kDa) are shown. Cupin-1.1 and Cupin-1.2 fibrils were used as control. **(c)** Birefringence of Vicilin fibrils stained with CR. Scale bar is equal to 20 μm. Left – transmitted light, right – polarized light. Cupin-1.2 fibrils were used as control. **(d)** Confocal microscopy of the fibrils stained with ThT. Fluorescence images of the ThT (top row), transmitted light images (middle row) and overlay (bottom row) are presented. The scale bar is equal to 15 μm. **(e)** Normalized to unity at the maximum absorption, fluorescence excitation and emission spectra and fluorescence decay curves of the ThT bound to Vicilin aggregates. The insets of the corresponding panels show the position of the maxima of the absorption and fluorescence excitation spectra, the values of the fluorescence quantum yield and lifetime of the bound to fibrils dye. By the dotted lines the long-wavelength regions of normalized absorption spectra (considering differences in the concentration of the bound dye) are presented. Decoding used colors is given in the Figure.

We have studied the resistance of the obtained by ‘seeding’ Vicilin fibrils to treatment with ionic detergents and have found that they are resistant to cold and hot SDS (Figure 3b). Staining of Vicilin fibrils with Congo red has demonstrated the clear apple-green birefringence indicating their amyloid structure (Figure 3c). Effective staining of the Vicilin fibrils by ThT (Figure 3d), a significant increase in the fluorescence quantum yield and lifetime of the bound dye (Figure 3e, right panels) and the presence of a pronounced long-wavelength shoulder in its absorption and fluorescence excitation spectra (Figure 3e, left panels) have confirmed the amyloid nature of these fibrils. It should be noted that along with the appearance of the long-wavelength shoulder in the absorption spectra of the bound dye, the concentration and, hence, the optical density of the bound ThT have increased (Figure 3e, dotted lines). This has led to the distortion of the short-wavelength region of the fluorescence spectra of ThT bound to Vicilin fibrils prepared by using the ‘seeds’. This is due to the secondary inner filter effect, which occurs when the absorption and fluorescence spectra of the sample substantially overlap and manifests in the reabsorption of light emitted by the sample. While determining the fluorescence quantum yield of the bound to fibrils ThT, this effect has been corrected by the usage of a specially developed experimental procedure^25^. Considering obtained results we have concluded that fibrillogenesis of the full-length Vicilin can be efficiently induced by the ‘seeds’ of pre-incubated Vicilin or Cupin-1.2 aggregates.

Taken together, we show that full-length Vicilin and both its domains Cupin-1.1 and Cupin-1.2 under certain conditions can form *bona fide* amyloid fibrils *in vitro*. Whereas Cupin-1.1 and Cupin-1.2 domains form amyloid fibrils after a simple incubation, amyloid fibril formation by Vicilin is induced by the addition of ‘seeds’ consisting of preformed aggregates of Vicilin or Cupin-1.2 but not Cupin-1.1.

### Vicilin, Cupin-1.1 and Cupin-1.2 exhibit amyloid properties being secreted to the surface of *Escherichia coli* cells and aggregate in *Saccharomyces cerevisiae*

We tested aggregation properties of Vicilin and its Cupin domains heterologously expressed *in vivo.* Firstly, we have analyzed the fibrillation of the Vicilin, Cupin-1.1 and Cupin-1.2 when these proteins are secreted to the surface of the *E. coli* cells in the C-DAG (Curli-Dependent Amyloid Generator) system^34^. We have found that the secretion of Vicilin and Cupin-1.1 has caused a vivid orange-red color of bacterial colonies on the plates with Congo red suggesting the extracellular aggregate formation by these proteins (Figure 4a). TEM has asserted this possibility demonstrating fibril formation by these proteins (Figure 4d). Interestingly, Cupin-1.2 has formed curved fibrils, while fibrils of Vicilin and Cupin-1.1 have been straighter (Figure 4d) correlating with a brighter color of the colonies secreting Vicilin and Cupin-1.1 and pale color of Cupin-1.2 colonies on the Congo plates (Figure 4.a). The polarization microscopy study of bacterial colonies secreting Vicilin, Cupin-1.1 and Cupin-1.2 fibrils has revealed a strong birefringence of all three variants confirming their amyloid properties (Figure 4b,c).

**Figure 4.**
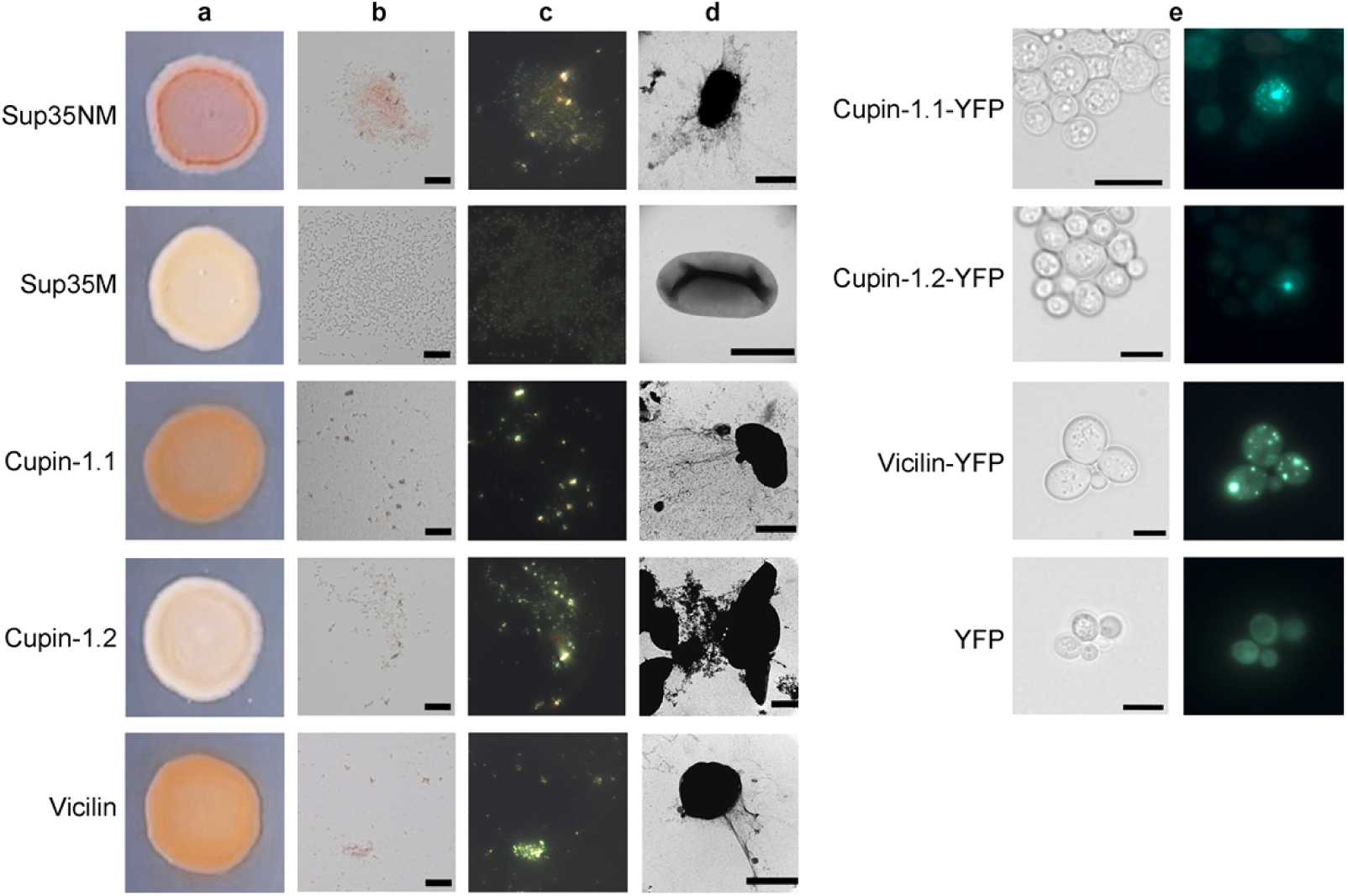
Amyloid properties of Vicilin and its Cupin domains produced in bacteria and yeast cells. **(a)** Congo red plate with *E. coli* cells secreting Vicilin, Cupin-1.1 and Cupin-1.2. The cells secreting either Sup35NM (amyloid) or Sup35M (soluble) proteins were used as the positive and negative controls, respectively. **(b, c)** Vicilin, Cupin-1.1 and Cupin-1.2 fibrils secreted by *E. coli* cells bind Congo Red and exhibit birefringence in polarized light. **(b)** Transmitted light and **(c)** polarized light images are shown, magnification 800x was used. **(d)** TEM images of the Vicilin, Cupin-1.1 and Cupin-1.2 fibrils secreted by *E. coli* cells. Scale bars are equal to 500 nm (all images) or 1 μm (Cupin-1.2). **(e)** The Vicilin, Cupin-1.1 and Cupin-1.2 proteins fused with YFP aggregate in the *S. cerevisiae* cells. Transmitted light (left) and fluorescent light (right) images are shown. Scale bar is equal to 5 μm.

In addition to the extracellular secretion of Vicilin, Cupin-1.1 and Cupin-1.2 in *E. coli*, we have overproduced these proteins fused with Yellow Fluorescent Protein (YFP) in yeast *S. cerevisiae* to test its intracellular aggregation. The results of this experiment have demonstrated that all three proteins, Vicilin, Cupin-1.1 and Cupin-1.2, fused with YFP have formed fluorescent aggregates in yeast cells (Figure 4e). Thus, we have found that full-length Vicilin and both its Cupin-1 domains demonstrate amyloid properties not only *in vitro* but being heterologously expressed *in vivo.*

### Vicilin forms amyloid aggregates in mature seeds *in vivo* and these aggregates resist canning and digestion by gastrointestinal enzymes

The most intriguing question was whether Vicilin forms amyloid aggregates in pea seeds *in vivo.* At the previous steps of this work we demonstrated the formation of detergent resistant Vicilin polymers in pea seeds by proteomic PSIA-LC-MALDI assay and its amyloid properties *in vitro* and heterologously expressed *in vivo*. To understand whether amyloid form of Vicilin can be detected in seeds histologically, we have analyzed the colocalization of Vicilin with amyloid-specific dye ThT on the cryosections of pea seed cotyledons (we used 30 days after pollination seeds where the accumulation of detergent-resistant form of Vicilin was detected by proteomic assay, Table 1). Immunofluorescent microscopy carried on unfixed cryosections has shown an almost total overlapping of the anti-Vicilin antibody and ThT signals indicating the presence of the Vicilin amyloid aggregates (Figure 5a-c). The anti-Vicilin antibody signal has been localized predominantly in the central vacuole and has revealed the presence of little granular compartments that have been proposed to be protein bodies, the membrane compartments where Vicilin typically accumulates^35^. To confirm this observation, we have isolated protein bodies using sucrose cushion sedimentation assay (see Materials and Methods) and have analyzed their staining with anti-Vicilin antibody and ThT. We have found that unfixed, air-dried protein bodies exhibit total overlapping of the anti-Vicilin and ThT signals suggesting that the amyloid Vicilin aggregates are located in protein bodies (laser scanning confocal microscopy, Figure 5d-f, and fluorescent microscopy, Figure S2). Finally, we have stained protein bodies with CR and have demonstrated that protein bodies are CR-positive and exhibit apple-green birefringence confirming their amyloid properties (Figure 5g).

**Figure 5.**
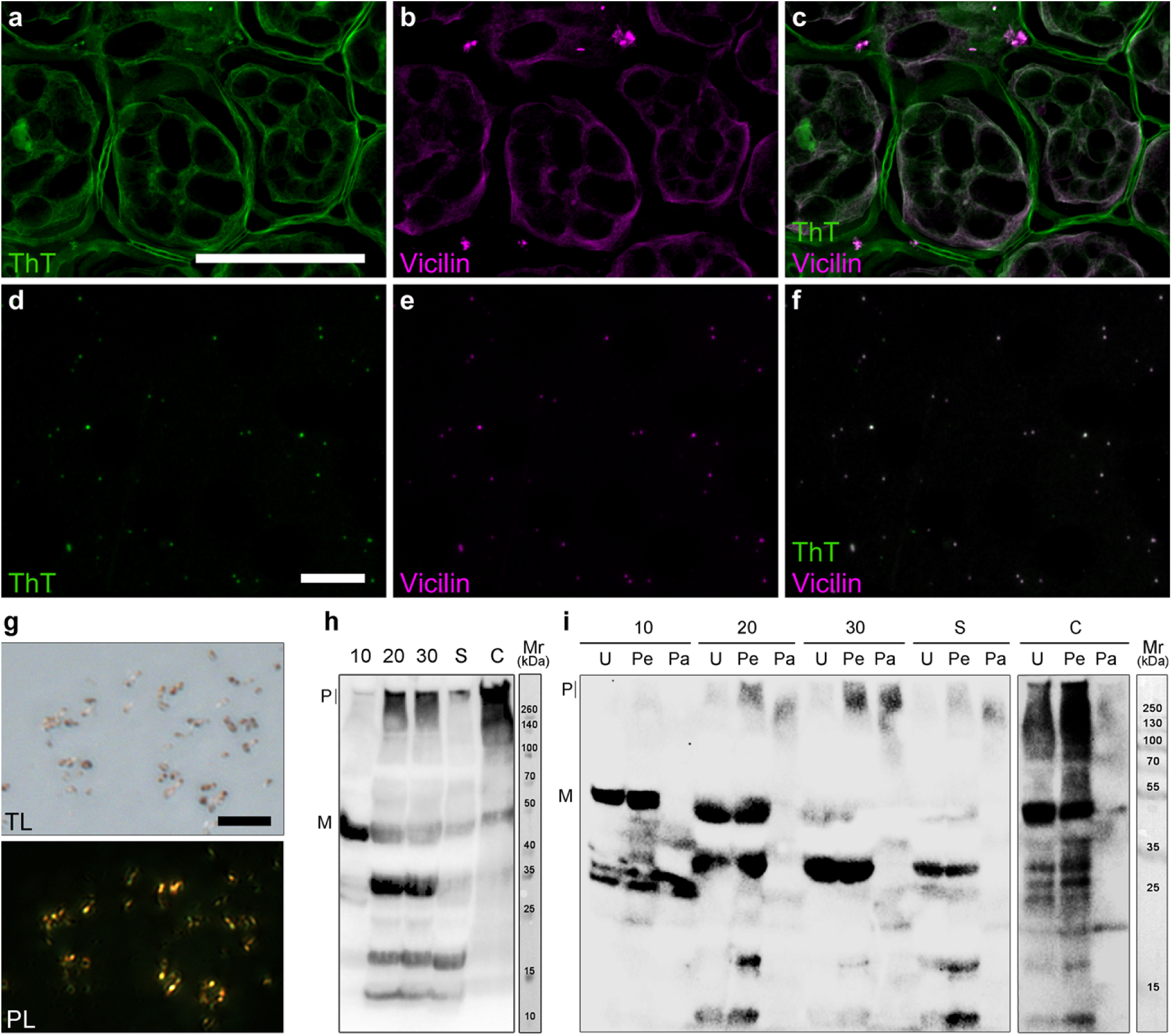
Vicilin forms amyloid aggregates *in vivo*. **(a-c)** *In situ* hybridization of anti-Vicilin antibodies with ThT on pea seed cryosections. Left image ThT channel of fluorescence (green), center – anti-Vicilin antibody (magenta), right – overlay (grey). **(d-f)** *In situ* hybridization of anti-Vicilin antibodies with ThT on protein bodies extracted from pea seeds. Scale bar is equal to 100 µm. **(g)** CR staining of isolated protein bodies. TL – transmitted light, PL – polarized light. Scale bar is equal to 20 µm. **(h)** Detergent-resistance of Vicilin aggregates isolated from pea seeds on different stages of maturation (10, 20, 30 days after pollination), germination (S, sprouts) and commercial canned peas (C, canned). **(i)** Protease resistance of Vicilin amyloids from pea seeds. U – untreated samples, Pe – treated with pepsin, Pa – treated with pepsin and then pancreatin, P – polymers, M – monomers. All samples in sections g-h were loaded onto the gel after 5 min boiling in buffer with 2 % SDS. Corresponding molecular weights are shown (kDa).

Next, we have studied the accumulation of the amyloid, detergent-resistant Vicilin aggregates in pea seeds using total protein lysates obtained at the different stages of the maturation or germination. Protein lysates were treated with 0.5 % SDS and 0.5 Tween20 for 10 min at RT, boiled for 10 min in the sample buffer containing 2% SDS (final concentration) and subjected to SDS-PAGE followed by western-blot with polyclonal anti-Vicilin antibody. The results of this experiment have confirmed aforementioned proteomic (Table 1) and histological (Figure 5a-g) data and demonstrated that amyloid aggregates of Vicilin are present in pea seeds (Figure 5h). We have found that these aggregates tend to accumulate during the seed maturation (from 10 to 30 days after pollination) and reach their maximum in mature seeds (Figure 5h). Notably, these aggregates rapidly disassemble in germinating seeds resulting in the formation of robust proteolytic band (Figure 5h). Since aggregates of Vicilin are highly stable, we have decided to analyze their presence in the commercial canned peas produced by ‘Bonduelle’ (Bonduelle Group, France) and ‘Heinz’ (H.J. Heinz, USA). The results of this experiment have shown that Vicilin amyloids persist canning and retain in these food products (Figure 5h, Figure S3).

We have analyzed the resistance of Vicilin amyloids to treatment with proteases. For this purpose, we have used *in vitro* protein digestibility assay (IVPD) that imitates gastrointestinal protein digestion^36^. Total protein lysates have been isolated from pea seeds, sprout cotyledons and canned peas. Next, lysates have been consequently treated with pepsin and pancreatin, boiled in the sample buffer with 2% SDS and subjected to SDS-PAGE and western-blot with polyclonal anti-Vicilin antibody. Results of this experiment have demonstrated that in contrast to monomers, Vicilin amyloids from pea seeds resist proteolytic digestion demonstrating high stability (Figure 5i). Notably, *in vitro* obtained Vicilin fibrils were not resistant to pepsin and pancreatin (Figure S4a); nevertheless, they demonstrated resistance to trypsin treatment (Figure S5), prolonged boiling with 2% SDS for 1 h (Figure S4b) and completely dissolved only by concentrated formic acid (Figure S6).

Taken together, we have demonstrated that Vicilin amyloids are present in pea seeds *in vivo* accumulating during the seed maturation, disassembling after germination and retaining in food products like canned peas.

### Vicilin amyloid fibrils exhibit toxicity for yeast and mammalian cells

Since we have found Vicilin forms amyloids both *in vivo* and *in vitro*, we have decided to analyze the effects of Vicilin amyloid formation. Vicilins are known to have carbohydrate-binding lectin activity resulting in their toxicity for fungi and exhibit functional dualism being not only storage but also pathogen-defense proteins^37^. In this study we have found that Vicilin can form at least two types of amyloid aggregates *in vitro*: less structured ones that are formed at the initial point of incubation and fibrils (Figures 1-3). To discriminate their effects, we tested toxicity of Vicilin, Cupin-1.1 and Cupin-1.2 non-fibrillar aggregates and fibrils for yeast cells at the same concentrations. The data obtained has demonstrated that Vicilin fibrils were very toxic for yeast culture resulting in significant yeast cells growth reduction after 48 h incubation (Figure 6a). Cupin-1.2 fibrils were extremely toxic resulting in almost complete death of cells (Figure 6a). In contrast to fibrils, non-fibrillar aggregates of Vicilin and Cupin-1.2 did not exhibit toxicity, while in the case of Cupin-1.1 both non-fibrillar aggregates and fibrils were toxic (Figure 6.a).

**Figure 6.**
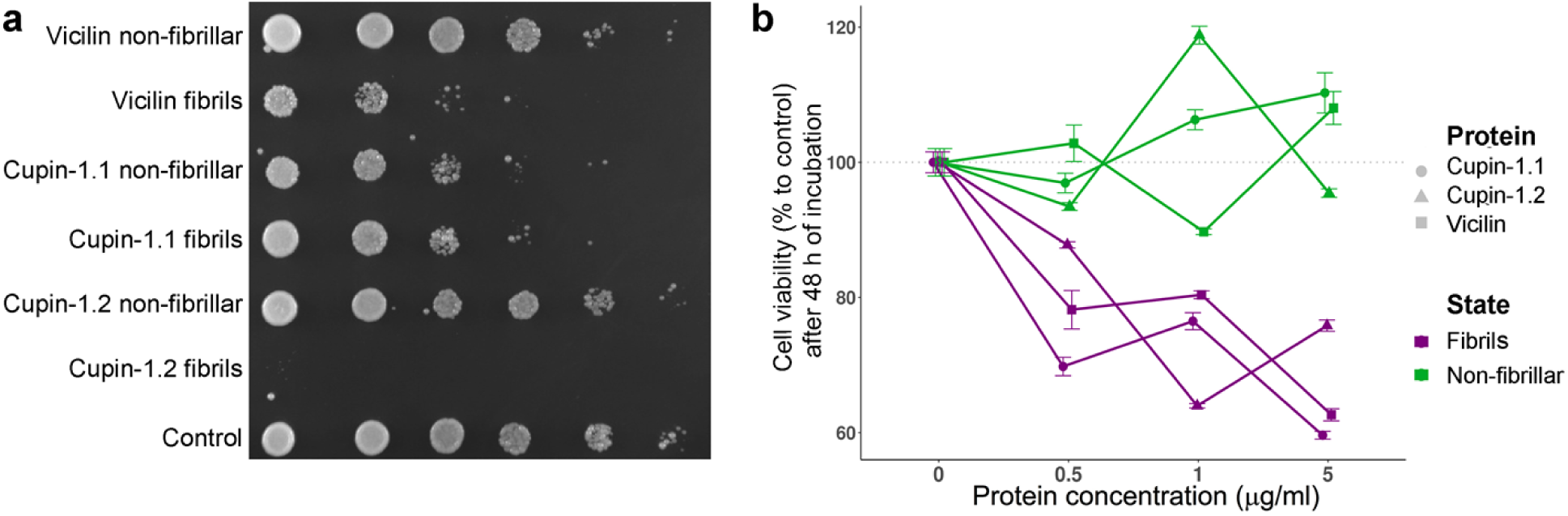
Vicilin fibrils are toxic for yeast and mammalian cells. **(a)** Analysis of the Vicilin, Cupin-1.1 and Cupin-1.2 fibrils and non-fibrillar aggregates toxicity for yeast cells. A series of tenfold dilutions of the liquid yeast culture is shown. Picture was taken after 48 h of incubation at 30° C. **(b)** Analysis of the Vicilin, Cupin-1.1 and Cupin-1.2 fibrils and non-fibrillar aggregates toxicity for mammalian cells. Dependence of the cell viability on the aggregates concentration is shown. Data was obtained after 48 h of incubation. Protein concentrations (µg/ml) are indicated. Scale bars correspond to the standard errors of the means.

To evaluate potential toxicity of Vicilin amyloids towards mammalian cells, we exposed human colorectal adenocarcinoma DLD1 cells to non-fibrillar aggregates and fibrils of Vicilin, Cupin-1.1 and Cupin-1.2. We did not observe any toxicity when DLD1 cells were treated with non-fibrillar aggregates of Vicilin, Cupin-1.1 and Cupin-1.2 (Figure 6b). On the contrary, Vicilin, Cupin-1.1 and Cupin-1.2 fibrils decreased viability of the cells in concentration dependent mode (Figure 6b).

Overall, Vicilin toxicity for fungal and mammalian cells depends on its structural characteristics and significantly increases when it forms amyloid fibrils. Both, Cupin-1.1 and Cupin-1.2 contribute to the toxicity but Cupin-1.2 fibrils exhibit especially high toxic effects for yeast cells.

## DISCUSSION

Plant seed maturation and transition to dormancy are accompanied by the desiccation that poses a stressful condition to all the cell components. Genome stability during desiccation is maintained via expression of genes related to the DNA repair and chromatin remodeling^38,39^ as well as chromatin compaction^40^. In contrast to the regulation of genome stability in seeds, molecular mechanisms underlying preservation of the seed proteins are poorly studied. Here, for the first time we have obtained an experimental proof that full-length plant seed storage 7S globulin Vicilin exhibits amyloid properties *in vivo* and *in vitro* (Figures 2-5). Amyloid aggregates of Vicilin accumulate in protein bodies during seed maturation and disassemble upon germination (Figure 5). These aggregates are protease- and detergent-resistant and bind amyloid-specific dyes ThT and Congo Red indicating their amyloid nature (Figures 2-5). Vicilin amyloids are efficiently solubilized only by concentrated formic acid being biochemically similar to extremely stable pathological human huntingtin exon-1 amyloids^41^. Such stability of the full-length protein is likely to be provided by the presence of two β-barrel domains Cupin −1.1 and −1.2 since they also form *bona fide* amyloid fibrils (Figures 1-2). Interestingly, structurally similar β-barrel domain-containing proteins that belong to Omp (outer membrane protein) superfamily of *Proteobacteria* also form amyloids^42,43^. Thus, several eukaryotic and prokaryotic proteins containing β-barrel domains are amyloid-forming and these domains are likely to be important amyloidogenic determinants.

Formation of amyloids by Vicilin *in vivo* corresponds to the fact that plants should develop specific mechanisms to rescue nutrient reservoir in seeds for growing embryos during long-term dehydration and unfavorable conditions. A series of studies of the seed storage proteins in various plants species has demonstrated that most of them including 7S Globulins, which Vicilin belongs to, form tri- or hexamers *in vivo^44^*, while isolated seed protein mixtures or solitary seed proteins tend to form amyloid-like fibrils *in vitro* after limited hydrolysis, at high temperatures or extremal pH^15^. Such amyloidogenic nature of seed storage globulins containing ancient β-barrel domains of Cupin superfamily is predicted bioinformatically in most land plant species suggesting for high evolutionary conservation of this feature^17^. Amyloids are probably most stable protein structures that resist different treatments and can persist in external environment for decades^45^. Thus, amyloid formation seems to be reasonable as evolutionary adaptation to provide a long-term survival of plant seeds. Moreover, similar examples of amyloids involved in protein storage were found in other kingdoms of life. For instance, egg envelope proteins of annual fish *Austrofundulus limnaeus* form amyloids during seasonal drought thus preventing dehydration^46^, while human peptide hormones accumulate as protein granules with amyloid structure that rapidly disintegrate and release functional monomeric hormones after pH change^47^. Thus, the storage function of amyloids is conservative not only in plants but exhibits cross-kingdom conservation between plants and animals. Significant decrease in the amounts of Vicilin amyloids, almost complete absence of the full-length Vicilin monomers and formation of proteolytic bands in germinating seeds (Figure 5) suggest that Vicilin amyloids are disassembled proteolytically, probably by serine and cystein proteases^48^, rather than by chaperone machinery.

Not only storage but also defense from pathogens (primarily, fungi and insects) function is typical for Vicilins^49^ and associated with lectin (carbohydrate-binding) properties of these proteins^50,51^ and their oligomerization^52,53^. Several lectins are known to form amyloid-like fibrils *in vitro* affecting their activity^54,55^. We have found out that the formation of amyloid fibrils significantly increases the toxicity of the full-length Vicilin for yeast in comparison with its unstructured aggregates (Figure 6) presupposing the role of amyloid formation in the defense function of this protein. Another property of lectins, in particular, Vicilins, is their allergenicity. Vicilin is one of the major plant-derived allergens, and its allergenic properties were found to be associated with its protease-resistant fragments^56,57^. Our finding that Vicilin forms amyloids *in vivo* explains its protease resistance (Figure 5) and suggests that these amyloids may represent major source of food allergy according to the data that *in vitro* generated lectin amyloids are phagocytized by macrophages and elicit an immune response^55^. Vicilin fibrils in high concentrations are toxic for mammalian cells (Figure 6) but concentrations of Vicilin amyloids in food products are many times lower, thus, their toxic effect for humans seems unlikely. Nevertheless, the presence of Vicilin amyloids in canned peas (Figure 5) can decrease food quality and increase allergic properties of seeds due to incomplete gastrointestinal digestion. Germination significantly reduces the amounts of amyloids (Figure 5), therefore, it is recommended to soak the seeds before eating to induce germination. Finally, the creation of novel plant varieties with decreased amyloid formation of storage proteins may represent a promising strategy for future agriculture to improve nutritional value and reduce allergenicity of plant seeds.

Overall, in this study we have identified plant protein that forms amyloids under native conditions *in vivo*, have found that these amyloids mediate protein storage in plant seeds and have demonstrated dynamics of their accumulation and disassembling, high stability, resistance to proteolytic digestion and canning as well as toxicity for fungal and mammalian cells.

## MATERIALS AND METHODS

### *P. sativum* L. *g*enetic lines and growing conditions

The pea (*P. sativum* L.) line Sprint-2 from the collection of ARRIAM (St. Petersburg, Russia) with determinate stem growth and early seed maturation was used^58^. To obtain seeds at various stages of maturation, pea seeds were sown in pots with ‘Terra vita Universal’ peat soil (MNPP FART Ltd., St. Petersburg, Russia). Plants were grown in a constant environment chamber (model VB 1514, Vötsch, Germany) at 16/8 h and 24/22°C day/night regime, 75% relative humidity, and around 10 000 lux illumination. The date of full opening of the flower was considered the date of pollination. Seeds intended for isolation of detergent-resistant protein fractions were collected 10, 20 and 30 days after pollination, which corresponded to the following stages of seed maturation: (i) flat pod, (ii) pod fill (green seeds) and (iii) yellow wrinkled pod^59^. Several plants cultivated were left until the dry harvest stage^59^ in order to study the effect of germination on the seed detergent-resistant fractions content afterwards. Harvested dry pea seeds were surface-disinfected for 6 min in 98% sulfuric acid, then thoroughly rinsed with sterile water and placed on Petri dishes containing sterile 1% agar-agar. After the seed germinating for 3 days at 27°C in the dark seedlings without signs of microbial contamination were selected and their cotyledons were cut off. Plant seeds and cotyledons collected were immediately frozen in liquid nitrogen and then stored at −80°C.

### Microbial strains, plasmids and cultivation conditions

The plasmids for analysis of aggregation of Vicilin, Cupin-1.1 and Cupin-1.2 fused with Yellow fluorescent protein (YFP) in yeast *S. cerevisiae* were obtained by the insertion of PCR amplified fragments of the corresponding genes amplified with primer pairs containing inserted *Bam*HI (reverse) and *Hind*III (forward) restriction sites (Table S2) and pea seed cDNA fragment as template into the pRS315-CUP1-SIS1-YFP plasmid^60^ by the *Bam*HI and *Hind*III sites, respectively. Total RNA from pea seeds was extracted using Trizol (Invitrogen, USA) and cDNA was prepared with SuperScript III reverse transcriptase (Invitrogen, USA). *E. coli* strain DH5α^61^ was used for plasmid amplification. The insertions of the corresponding genes were confirmed by the sequencing by using the CUP1 primer^60^.

The *S. cerevisiae* 1-OT56 strain (*MAT*a *ade1-14*_UGA_ *his3 leu2 trp1-289*_UAG_ *ura3* [*psi*^-^][*PIN*^+^])^60^ was transformed with constructed plasmids. Yeast cultivation was performed on selective agar media at 30°C for 4 days followed by replacing on agar media with 150 µL ml^-1^ of CuSO_4_ for *CUP1* promoter induction. Fluorescence was analyzed using a Zeiss Axio Imager A2-fluorescent microscope (Carl Zeiss, Germany).

To construct plasmids for Vicilin, Cupin-1.1, and Cupin-1.2 export by curli-dependent amyloid generator (C-DAG) system the fragments of interest were amplified by PCR using pairs of primers flanked with *Not*1 (forward) and *Sal*1 (reverse) restriction sites (Table S2) and pea seed cDNA as template. The PCR products and pExport vector^34^were digested by *Not*1 and *Sal*1 restriction enzymes and then ligated. Plasmids encoding yeast Sup35NM (aa 2–253) and Sup35M (aa 125–253) fused with CsgA signal sequence were constructed previously^34^.

### C-DAG assay

Analysis of amyloid properties of Vicilin, Cupin-1.1, and Cupin-1.2 with the usage of curli-dependent amyloid generator (C-DAG) system was performed as described earlier^34^. To export proteins on the cell surface *E. coli* strain VS39^34^ was transformed with pExport-based plasmids, encoding Vicilin, Cupin-1.1, and Cupin-1.2 fused with CsgA signal sequence. Export of amyloid-forming Sup35NM and non-amyloidogenic Sup35M proteins was used as positive and negative control of amyloid formation respectively. Birefringence analysis was performed with a usage of Axio Imager A2 transmitted light microscope (Zeiss) equipped with a 40x objective and cross-polarizers. For transmission electron microscopy analysis incubation on inducing plates without Congo red dye was used.

### Mammalian cell lines and toxicity assay

Human DLD1 cells were seeded at 5*10^4^ cells per well density on 24-well plates and exposed to protein samples analyzed. At indicated time points, medium was replaced with 0,5 mg/ml MTT solution for 1h at 37°C. Then formazan was dissolved in DMSO, and optical densities were measured at 572 nm wavelength using the Multiscan Ex spectrophotometer (Thermo, USA). Data were presented as the mean of three independent experiments ± the standard error of the mean.

### PSIA-LC-MALDI proteomic assay

The PSIA-LC-MALDI proteomic assay for the identification of protein complexes resistant to the treatment with ionic detergents has been published previously^22^. Here, we have used this protocol with several modifications. Frozen pea seed cotyledons (1 g) were disrupted with pestle and mortar in liquid nitrogen. Resulting powder was dissolved in 5 ml of PBS buffer and treated with 1% SDS and 0.2% Tween-20 detergents for 15 min. Next, lysate was sedimented at 10 000 g for 5 min, supernatant was moved to new tube and sedimented again. Resulting supernatant was loaded onto the top of 1.5 ml of 25% sucrose/PBS with 0.1% SDS cushion and centrifuged for 8 h, 225 000 g at 18°C in L8-70 ultracentrifuge (Bechman Coulter, USA), Type50 Ti angle rotor (Bechman Coulter, USA). Resulting pellets were washed in 5 ml of distilled water and centrifugated again (1.5 h, 225 000 g, 4°C, L8-70 ultracentrifuge (Bechman Coulter, USA), Type50 Ti angle rotor (Bechman Coulter, USA)). This step was repeated twice.

Next, samples were lyophilized, treated with formic acid to solubilize proteins and lyophilized again. Salts and detergents were removed and samples were subjected to trypsinolysis stage followed by reverse phase high performance liquid chromatography and mass-spectrometry as described earlier^22^ During analysis, preset parameters of ‘Mass tolerance’ were used (precursor mass tolerance 50 ppm, fragment mass tolerance 0.9 Da). Peptide Calibration Standard II 8222570 (Bruker Daltonics) was applied as a standard sample. Carboxymethylation of cysteine, partial oxidation of methionine, and one skipped trypsinolysis site were considered as permissible modifications. The obtained mass-spectra were matched to the corresponding proteins using NCBI database. The BioTools software (Bruker, Bremen, Germany) was used for manual validation of protein identification.

### Protein expression and purification

To express the Vicilin, Cupin1-1, Cupin1-2 terminally fused with a 6x-His tag, an Alicator kit (Thermo Scientific, USA) was used. The fragments were PCR-amplified using respective primer pairs (Table S2) and pea seed cDNA as template. The PCR-amplified fragments were inserted into the pAlicator vector (Thermo Scientific, USA) according to the manufacturer’s recommendations. The correctness of the resulting pAc-Vicilin, pAc-Cupin-1.1 and pAc-Cupin-1.2 plasmids was verified by sequencing with the primers provided by the manufacturer (Thermo Scientific, USA).

For protein expression, *E. coli* strain BL21 (New England Biolabs, USA) was used. The overproduction of recombinant proteins was carried out in 2TYa media supplemented with 0.1 mM IPTG. Cultures were grown at 37°C for 4 h. Proteins were purified in denaturing conditions (in the presence of 8M urea) according to a previously published protocol^62^ without the Q-sepharose purification step. A one-step purification procedure with a Ni-NTA agarose (Invitrogen, USA) column was performed according to the manufacturer’s recommendations. Proteins were concentrated using ethanol.

### *In vitro* protein fibrillation

For initiation of Vicilin and its domains aggregation *in vitro*, different buffers and incubation conditions were used. At the first stage the proteins in concentration 0.5 mg/ml were incubated in phosphate buffer (pH 7.4) at room temperature and at 50°C with constant stirring for 2 weeks. Furthermore, the proteins in the same concentration were dissolved in 50% HFIP (Sigma-Aldrich, USA) and incubated for 7 days. Afterwards, the HFIP was evaporated under a stream of nitrogen, and the samples were stirred for an additional 7 days. These conditions were also used for experiments with seeding. ‘Seeds’ that were prepared on the basis of pre-incubated Vicilin, Cupin-1.1 or Cupin-1.2 aggregates were added to the samples at the beginning of fibrillogenesis in 1% (v/v) concentration.

### Pea seed protein extraction, in gel separation and transfer

A standard protocol was used for total protein isolation from pea seeds. Three seeds collected at indicated stage of maturation or cotyledons of three seeds after germination were analyzed in each sample. Canned peas produced by ‘Bonduelle’ (Bonduelle Group, France), ‘Heinz’ (H.J. Heinz, USA) were purchased in a supermarket in St. Petersburg, Russia. Homogenization of seeds was performed using glass beads. Protein concentrations were equilibrated using Qubit 3 Fluorometer (Invitrogen, USA). Then detergents were added sequentially to the final concentrations: 0.5% Tween20 (Helicon, Russia), 0.5% SDS (Helicon, Russia) followed by 10 min incubation at RT. Further, if necessary, *In vitro* protein digestibility (IVPD) was applied. Next, samples were boiled in the presence of 2% SDS for 5 min and loaded onto the 10% SDS-PAGE gel (Bio-Rad, USA). After SDS-PAGE, wet transfer (Bio-Rad, USA) was performed using PVDF membrane (GE Healthcare, USA).

### *In vitro* protein digestibility (IVPD)

Pea seed proteins were digested *in vitro* as described previously^36^. After each step of enzyme treatment samples were collected and inactivated by heating at 100°C for 5 min. Pepsin (Roche, Germany) and pancreatin from porcine pancreas (Sigma, USA) were used. Samples were checked with SDS-PAGE followed by western-blot with polyclonal anti-Vicilin antibod? (Imtek, Russian Federation). All analyses were performed in quadruplicate.

### Immunochemical analysis

To obtain polyclonal anti-Vicilin antibodies from serum of a healthy rabbit, immunization with the antigen of the protein received according to the protocol described above was performed. Rabbit immunization and purification using affinity chromatography with rabbit antisera on a sorbent with immobilized recombinant Vicilin protein were carried out in Imtek company (Imtek, Russian Federation). Dilution 1:1 000 was used. Further goat anti-rabbit IgG (H+L) secondary antibody (Thermo Scientific, USA) was used in dilution 1:33 000. Also, 6x-His epitope tag antibody (4A12E4, Invitrogen, USA) was used in dilution 1:5 000, then goat anti-mouse IgG (H+L) secondary antibody (Invitrogen, USA) was used in dilution 1:5 000.

### Protein body isolation

Protein bodies were extracted from fresh pea seeds of 30 days after pollination according to modified protocol for *Zea mays* protein bodies^63^. In particular, 2 g of the seeds were grounded with cooled mortar and pistil in the presence of 5 ml of 10% (w/w) sucrose in phosphate buffer solution (PBS) containing 5 mM phenylmethylsulfonyl fluoride (PMSF). Resulting homogenate was filtered through the nylon cloth and filtrate was then centrifuged at 500 g and 4°C for 10 min. Supernatant was layered onto discontinuous sucrose gradient comprising of 1, 1.5 and 1.5 ml of 70%, 50% and 20% (w/w) sucrose cushions in PBS respectively, in the open-top thickwall polycarbonate tube (Bechman Coulter, USA) and centrifuged in L8-70 ultracentrifuge (Bechman Coulter, USA) in Type50 Ti angle rotor (Bechman Coulter, USA) at 36 500 g and 4°C for 1 h. Protein bodies were collected from the boundary of 70% and 50% cushions as a part of opaque white halo. Approximately 100 µl of this fraction was extracted, tenfold diluted with 10% sucrose in PBS and centrifuged at 12 000 g, 4°C,15 min. The supernatant was discarded and the pellet was resuspended in 10% sucrose in PBS.

### *In situ* hybridization and fluorescent microscopy

Slices preparation: mature pea seeds were glued to holder with O.C.T. (Tissue-Tek, Sakura Finetek, Japan) and cut with a cryostat (Leica CM3050S, Leica Biosystems Nussloch GmbH, Germany) at −16°C to 20-μm-thick sections which were mounted on poly-l-lysine coated slides. To promote better adhesion, slides were kept at least 12 h at 37°C. Protein bodies preparation: sucrose solution of protein bodies was applied on a poly-l-lysine coated slides and air-dried for 30 min at room temperature. Slides were washed twice for 5 min in TBS (tris-buffered saline, 0.1M), blocked for 2 h in humid chamber at room temperature with solution containing 10% normal goat serum (Sigma, USA), 0.1% Triton X-100 (Sigma, USA) on TBS. Sections were incubated with primary antibodies in a humid chamber at 4°C overnight. Polyclonal rabbit-anti-Vicilin antibodies were applied at concentration 1:100 in TBS with 0.01% Triton X-100 and 1% Normal goat serum. The primary antibodies were detected by 2 h incubation at 37°C with secondary antibodies solution: goat anti-rabbit IgG, conjugated to Alexa Fluor 568 (Life Technologies, USA) diluted 1:250 in TBS containing 1% normal goat serum and 0.01% Triton X-100. Sections were washed twice with TBS. Slides were additionally stained for 40 min with 0.5% solution of ThT (Thioflavin T, Sigma, USA) in 0.1N HCl, briefly differentiated in 70% ethanol, washed by TBS and mounted in 80% TBS-glycerol. Longer staining times were avoided due to rapid washing-off of immunofluorescent label. A control for nonspecific staining was performed by replacing the primary antibody produced in rabbit with 10% normal rabbit serum. Sections were analyzed with the fluorescence microscope Leica DM5500 and scanning confocal microscope Leica TCS SP5 (Leica, Germany).

### Transmission electron microscopy

Micrographs were obtained using a transmission electron microscope Libra 120 (Carl Zeiss, Germany). The samples were placed on nickel grids coated with formvar films (Electron Microscopy Sciences, USA). To obtain electron micrographs, the method of negative staining with a 1% aqueous solution of uranyl acetate was used.

### Polarization microscopy

Congo red staining of amyloid samples was performed using saturated Congo red (Sigma, USA) solution filtered through 45 µm filter (Millipore, USA). Samples stained with Congo red were air-dried and rigorously washed with distilled water. Birefringence was analyzed using Zeiss Axio Imager A2 (Carl Zeiss, Germany) polarization microscope equipped with cross polarizers.

### ThT staining and confocal microscopy

Thioflavin T (ThT) UltraPure Grade (AnaSpec, USA) without after-purification was used. ThT-fibrils tested solutions were prepared by equilibrium microdialysis using a Harvard Apparatus/Amika device (USA). Equilibrium microdialysis was performed with a concentration of aggregates ∼0.5 mg/ml and initial concentration of ThT ∼32 μM. Spectroscopic study of the sample and reference solutions prepared by proposed approach allowed us to determine the photophysical characteristics ThT bound to tested amyloids^24^.

For obtaining the fluorescence images of the ThT-stained fibrillar structures confocal laser scanning microscope Olympus FV 3000 (Olympus, Japan) was used. We used the oil immersion objective with a 60x magnification, numerical aperture NA 1.42 and laser with excitation line 405 nm.

### Spectral measurements

The absorption spectra of the samples were recorded using a U-3900H spectrophotometer (Hitachi, Japan). The absorption spectra of proteins aggregates and ThT in their presence were analyzed along with the light scattering using a standard procedure^64^. The concentrations of Vicilin, Cupin-1.1, Cupin-1.2 and ThT in solutions were determined using molar extinction coefficients of ε_280_ = 47271 M^-1^cm^-1^, ε_280_ = 16928 M^-1^cm^-1^, ε_280_ = 18549 M^-1^cm^-1^ and ε_412_ = 31589 M^-1^cm^-1^, respectively.

Fluorescence and fluorescence excitation spectra were measured using a Cary Eclipse spectrofluorimeter (Varian, Australia). Fluorescence of ThT was excited at a wavelength of 440 nm and registered at 490 nm. A PBS solution of ATTO-425, whose fluorescence and absorption spectra are similar to that of ThT, was taken as a reference for determining the fluorescence quantum yield of ThT bound to fibrils. The fluorescence quantum yield of ATTO-425 was taken as 0.9 (ATTO-TEC Catalogue 2009/2010 p.14). The spectral slits width was 5 nm in most of experiments. Changing the slit widths did not influence the experimental results. Recorded fluorescence intensity was corrected on the primary inner filter effect with the use of previously elaborated approach^25^.

CD spectra in the far UV-region were measured using a J-810 spectropolarimeter (Jasco, Japan). Spectra were recorded in a 0.1 cm cell from 260 to 200 nm. For all spectra, an average of three scans was obtained. The CD spectrum of the appropriate buffer was recorded and subtracted from the samples spectra. It has turned out that the recorded CD spectra have a clear distortion in the region of 250-260 nm (Figure 2a). According to literature light scattering of macromolecules can substantially distort the CD spectra (see, for example,^65^). We have shown that the degree of the observed distortion (difference of recorded values from 0) actually correlates with the turbidity of the samples, which, in turn, is determined by the size of the studied aggregates (Figure 2a, Inset). We attempted a quantitative analysis of the secondary structure by the CDPro program using three different regression methods (Selcon, Contin, and CDSSTR) and several basic sets of proteins with a known secondary structure (the sets include from 37 to 56 soluble, membrane, and denatured proteins with different content of the secondary structure). Since such results could be arbitrary because the standard basic sets of proteins used to estimation of the secondary structure content are not representative for the analysis of the spectra of protein aggregates, we have also conducted a visual analysis of the recorded spectra with the use of CD spectra of proteins and peptides with representative secondary structures^28^.

### Time-resolved fluorescence measurements

Fluorescence decay curves were recorded by a spectrometer FluoTime 300 (PicoQuant, Germany) with the Laser Diode Head LDH-C-440 (*λ*_*ex*_ = 440 nm). The fluorescence of ThT was registered at *λ*_*em*_ = 490 nm. The measured emission decays were fit to a multiexponential function using the standard convolute-and-compare nonlinear least-squares procedure^66^. In this method, the convolution of the model exponential function with the instrument response function (IRF) was compared to the experimental data until a satisfactory fit was obtained. The fitting routine was based on the nonlinear least-squares method. Minimization was performed according to Marquardt^67^.

### Data analysis

All experiments in this work were performed in at least three repeats. Data in figures is presented as the mean ± the standard error of the mean. In human cell toxicity assay, a dependence of the toxicity on the concentration was considered as significant if p-value of the regression coefficient in generalized linear model (glm function in R language) was lower than 0.05 and therefore the regression coefficient was significantly different from zero.

## ACKNOWLEDGEMENTS

This work was supported by Russian Science Foundation, grant 17-16-01100. The work of A.I. Sulatskaya (polymorphism of amyloid aggregates under various conditions) was awarded by RF President Fellowship SP-841.2018.4.

## AUTHOR CONTRIBUTIONS

Conceptualization: AAN; performed experiments: KSA, MVB, AIS, MEB, AOK, MIS, OYS, EAA, PAZ, ANL, KVV, YVM, EYK and AAN; wrote original draft: KSA, YVM, MVB, AIS, MIS, OYS, EAA, PAZ, ANL, OND and AAN; analyzed results: KSA, MVB, AIS, MEB, AOK, MIS, OYS, EAA, PAZ, ANL, KVV, YVM, IMK, KKT, IAT, EYK, OND and AAN; discussed results: KSA, MVB, AIS, MEB, AOK, MIS, OYS, EAA, PAZ, ANL,

KVV, YVM, IMK, KKT, IAT and AAN; funding acquisition: AAN.

